# Cofactor specificity of glucose-6-phosphate dehydrogenase isozymes in *Pseudomonas putida* reveals a general principle underlying glycolytic strategies in bacteria

**DOI:** 10.1101/2021.01.08.426012

**Authors:** Daniel C. Volke, Karel Olavarría, Pablo Ivan Nikel

## Abstract

Glucose-6-phosphate dehydrogenase (G6PDH) is widely distributed in nature and catalyzes the first committing step in the oxidative branch of the pentose phosphate (PP) pathway, feeding either the reductive PP or the Entner-Doudoroff pathway. Besides its role in central carbon metabolism, this dehydrogenase also provides reduced cofactors, thereby affecting redox balance. Although G6PDH is typically considered to display specificity towards nicotinamide adenine dinucleotide phosphate (NADP^+^), some variants accept nicotinamide NAD^+^ similarly (or even preferentially). Furthermore, the number of G6PDH isozymes encoded in bacterial genomes varies from none to more than four orthologues. On this background, we systematically analyzed the interplay of the three G6PDH isoforms of the soil bacterium *Pseudomonas putida* KT2440 from a genomic, genetic and biochemical perspective. *P. putida* represents an ideal model to tackle this endeavor, as its genome encodes numerous gene orthologues for most dehydrogenases in central carbon metabolism. We show that the three G6PDHs of strain KT2440 have different cofactor specificities, and that the isoforms encoded by *zwfA* and *zwfB* carry most of the activity, acting as metabolic ‘gatekeepers’ for carbon sources that enter at different nodes of the biochemical network. Moreover, we demonstrate how multiplication of G6PDH isoforms is a widespread strategy in bacteria, correlating with the presence of an incomplete Embden-Meyerhof-Parnas pathway. Multiplication of G6PDH isoforms in these species goes hand-in-hand with low NADP^+^ affinity at least in one G6PDH isozyme. We propose that gene duplication and relaxation in cofactor specificity is an evolutionary strategy towards balancing the relative production of NADPH and NADH.

**Importance:** Protein families have likely arisen during evolution by gene duplication and divergence followed by *neo*-functionalization. While this phenomenon is well documented for catabolic activities (typical of environmental bacteria that colonize highly polluted niches), the co-existence of multiple isozymes in central carbon catabolism remains relatively unexplored. We have adopted the metabolically-versatile soil bacterium *Pseudomonas putida* KT2440 as a model to interrogate the physiological and evolutionary significance of co-existing glucose-6-phosphate dehydrogenase (G6PDH) isozymes. Our results show that each of the three G6PDHs encoded in this bacterium display distinct biochemical properties, especially at the level of cofactor preference, impacting bacterial physiology in a carbon source-dependent fashion. Furthermore, the presence of multiple G6PDHs differing in NAD^+^- or NADP^+^-specificity in bacterial species strongly correlates with their predominant metabolic lifestyle. Our findings support the notion that multiplication of genes encoding cofactor-dependent dehydrogenases is a general evolutionary strategy towards achieving redox balance according to the growth conditions.

## Introduction

Glycolysis, the set of reactions that converts glucose and other sugars into biomass building blocks, is a ubiquitous metabolic feature across the tree of life. Glycolysis is often equated to the Embden-Meyerhof-Parnas (EMP) pathway, as it was firstly described in eukaryotes and the model Gram-negative bacterium *Escherichia coli*. However, glycolytic strategies are rather diverse, and several biochemical architectures (ranging from linear to highly-interconnected cyclic networks) are deployed to metabolize glucose in different microbial species. Flamholz et al. (1) pointed that there is no ‘superior’ glycolysis (in terms of overall efficiency), and that each metabolic architecture can potentially confer survival advantages under certain environmental conditions. For instance, the Entner-Doudoroff (ED) pathway (and modified versions thereof) is a glycolytic strategy often found besides, instead of, or in combination with the EMP pathway (2–4). The ED pathway produces one ATP less per glucose than the EMP pathway (5), but it requires considerably less enzymatic resources (i.e. it is considered to be more efficient in terms of protein allocation). The ED pathway seems to prevail in bacteria with an aerobic lifestyle, where a large ATP fraction is generated by aerobic respiration instead of substrate-level phosphorylation (1). Besides providing precursors for biomass build-up, glycolytic routes recycle essential energy and redox cofactors—i.e. adenosine triphosphate (ATP), reduced nicotinamide adenine dinucleotide (NADH) and reduced nicotinamide adenine dinucleotide phosphate (NADPH). While ATP is mostly used as an energy currency, NADH and NADPH are used as carriers of reducing equivalents. Even though the chemical structures of NADH and NADPH are very similar, the NADH/NAD^+^ and NADPH/NADP^+^ redox ratios are very different—mirroring the different functions of these cofactors in the cell (6). NADH is primarily re-oxidized in the respiratory chain to generate ATP, while NADPH drives anabolic reactions (7) and is also used to reduce glutathione as a first line of defense to oxidative stress (8).

Strategies to adequately balance consumption and production of NADH and NADPH under different environmental conditions are a crucial trait for every cell. Besides altered cofactor specificities in dehydrogenases of central carbon metabolism, allosteric regulation mechanisms and rapid adjustment of carbon fluxes within the metabolic network are typically observed across bacterial species (9, 10). An important reaction for cofactor balance is catalyzed by glucose-6-phosphate dehydrogenase (G6PDH), the first step of the oxidative pentose phosphate (PP) pathway and the ED pathway. In the *E. coli*-centric (and prevailing) view of metabolism, this reaction is thought to serve as the primary source of NADPH (10–12) and the EMP pathway is the predominant catabolic route for hexoses in *E. coli*. Building on this notion, when the cofactor specificity of G6PDH of a given organism has not been characterized in detail, it is often assumed that it will prefer NADP^+^. However, several lines of evidence indicate that this is not the case for bacteria beyond *E. coli* (13, 14). For example, when this linear route is blocked—*e.g.* in a Δ*pgi* mutant—and all the glycolytic flux is forced through the reaction catalyzed by G6PDH, the strict cofactor preference of the G6PDH from *E. coli* causes an NADPH imbalance, leading to severe effects on bacterial growth (15, 16). In that case, mechanisms that prevent NADPH accumulation—*e.g.* over-expressing *sthA*, which encodes for a soluble transhydrogenase (15), or substituting the native G6PDH by an NAD^+^-preferring orthologue (16)—partially rescues the growth rate of the Δ*pgi* mutant in cultures using glucose as the sole carbon source. This state of affairs prompts the following questions: what is the cofactor specificity of G6PDH in bacteria relying on the ED pathway for glycolysis, where most of the glycolytic flux is channeled through this dehydrogenase, and how do these bacteria adjust NADPH balances? Which is the role of G6PDH in the NAD(P)H balancing mechanisms in those organisms? Studying the properties of G6PDHs in a model organism where these questions can be addressed is thus critical to understand the underlying regulatory mechanisms.

The use of the ED pathway is widespread in *Pseudomonas*, and previous studies reported on the presence of several copies of G6PDH-encoding genes in this genus (17–20). *Pseudomonas putida* KT2440 and other members of this genus use a cyclic glycolysis **(Fig. 1)**, employing the ED and PP pathways in combination with gluconeogenesis through the EMP route to oxidize hexoses (21). This structure, termed EDEMP cycle, was described in *P. putida* KT2440 as a biochemical mechanism that adjusts NADPH production levels through cycling the flux through the upper glycolysis (21, 22). Besides this main set of reactions in the cytoplasm, *P. putida* harbors periplasmic enzymes for the direct oxidation of glucose to gluconate. Previous studies indicated that around 80% of glucose is taken up as gluconate or 2-ketogluconate, while the remaining 20% is phosphorylated to glucose 6-phosphate (21, 23). G6PDH is a central enzymatic step for glycolysis in *P. putida*, and the three isoforms of this enzyme are encoded by *zwfA* (*PP_1022*), *zwfB* (*PP_4042*) and *zwfC* (*PP_5351*). G6PDH-A, encoded by *zwfA*, was shown to be highly produced under glycolytic conditions, with the transcription of the cognate gene (negatively) regulated by HexR (24). However, the function (if any) of *zwfB* and *zwfC* and their corresponding products, G6PDH-B and G6PDH-C, remained elusive thus far.

**Figure 1.**
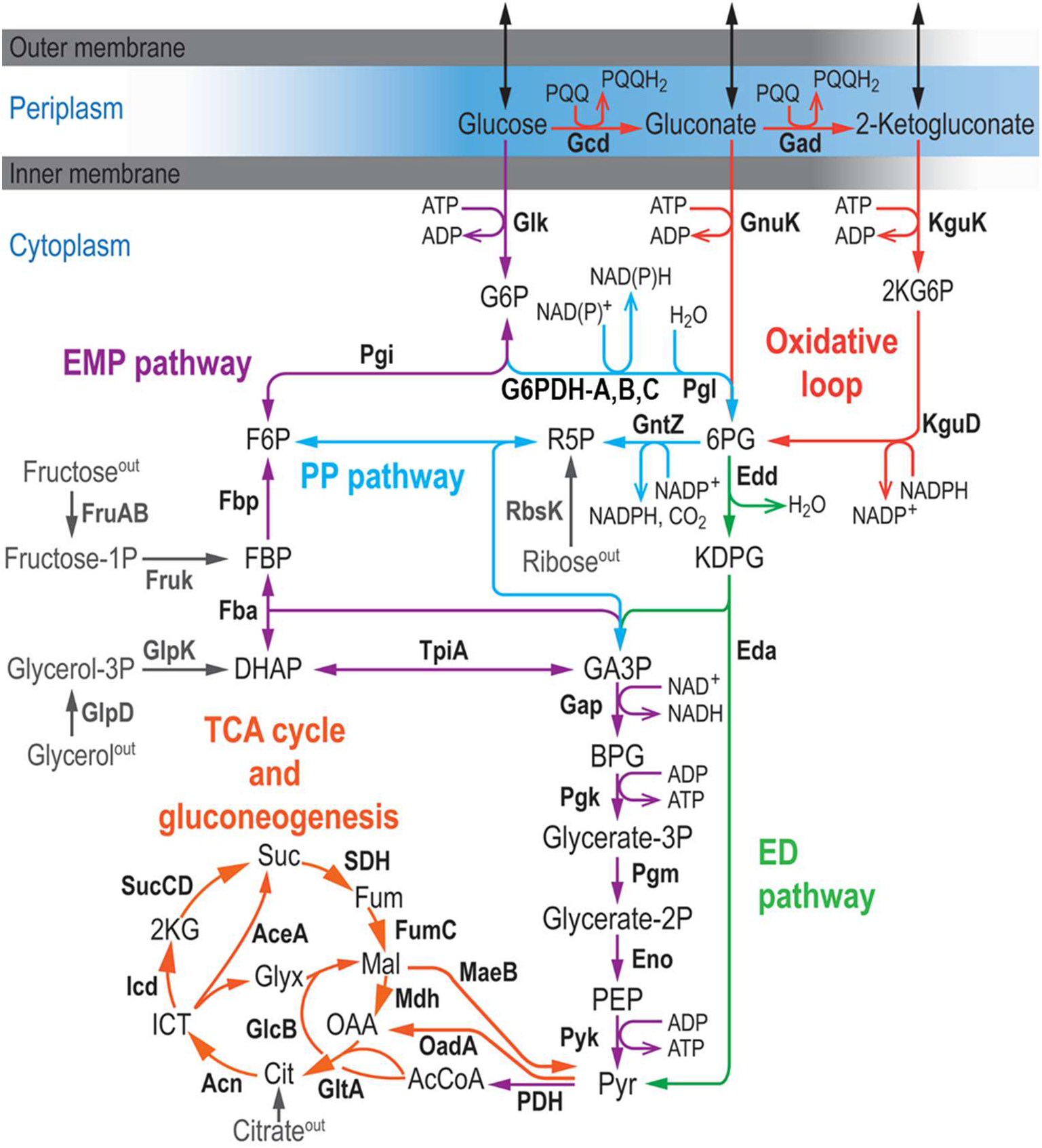
Organization of central carbon metabolism in *P. putida* KT2440. The metabolic network is sketched in six modules, depicted in different colors. Glucose is partly oxidized in the periplasm to yield gluconate and 2-ketogluconate, but both the sugar and its oxidized derivatives can be phosphorylated in the cytoplasm and metabolized through the Entner-Doudoroff (ED) pathway. Part of the carbon flux is cycled through the (incomplete) Embden-Meyerhof-Parnas (EMP) pathway operating in a gluconeogenic fashion, and a fraction of the glycolytic flux is channeled through the pentose phosphate (PP) pathway. The connecting point between the ED and EMP routes is the reaction catalyzed by glucose-6-phosphate dehydrogenases (G6PDH), encoded by three genes in strain KT2440: *zwfA*, *zwfB* and *zwfC.* Furthermore, the upper domain glycolysis is connected to the tricarboxylic acid (TCA) cycle *via* the lower EMP pathway. For simplicity, some reactions are either lumped together or not shown in the diagram, and entry points of alternative carbon sources (i.e. fructose, ribose, glycerol and citrate) are indicated in grey. Abbreviations for the key metabolites and intermediates in the network are as follows: 2K6PG, 2-keto-6-phosphogluconate; KDPG, 2-keto-3-deoxy-6-phosphogluconate; 6PG, 6-phosphogluconate; DHAP, dihydroxyacetone phosphate; FBP, fructose-1,6,-bisphosphate; GA3P, glyceraldehyde-3-phosphate; R5P, ribose-5-phosphate; BPG, 1,3-bisphosphoglycerate; PEP, phosphoenolpyruvate; Pyr, pyruvate; AcCoA, acetyl-coenzyme A; Cit, citrate; ICT, isocitrate; 2KG, 2-ketoglutarate; OAA, oxaloacetate; Suc, succinate; Fum, fumarate; Mal, malate; and Glyx, glyoxylate.

Given such characteristics, we have adopted this bacterium as a model to explore the role of the different G6PDHs. We show that *P. putida* relies on the activity of isozymes with different cofactor specificity for the catabolism of several carbon sources. These enzyme variants were found to be especially important for growth on fructose and the pentose ribose, as nearly all the carbon flux goes through the G6PDH reaction. We also show that the presence of dual-cofactor and multiple G6PDHs isozymes are a metabolic signature that strongly correlates with the use of the ED pathway as the main glycolytic strategy. As exposed by sequence analysis of G6PDHs across more than thousand bacterial species, this metabolic strategy appears to be widespread in the bacterial kingdom.

## Results

### Growth patterns of single, double and triple zwf mutants of P. putida KT2440 expose the physiological role of each isoform

In order to shed light on the roles of the G6PDH isozymes, *zwfA*, *zwfB* and *zwfC* were deleted in *P. putida* KT2440. The eight possible combinations of gene deletions comprising these three genes were constructed towards revealing the individual contribution of each variant to the overall physiology of the cells. These single and multiple *zwf* mutants were generated by marker-less deletion based on homologous recombination (25, 26), and thoroughly checked for correctness by PCR amplification of the relevant loci and DNA sequencing **(Fig. S1)**.

In order to test the coarse phenotype of the resulting strains, the mutants were first grown under a number of cultivation conditions and the growth rates were determined. All mutant strains had the same specific growth rate (μ) when grown in complex lysogeny broth (LB) medium, indicating that mutations in *zwf* did not affect significantly cellular mechanisms involved in anabolism or bacterial division. Physiological characterization was also done in M9 minimal medium supplemented with different carbon sources **(Fig. 2A)**, selected according to the different catabolic modules involved in their processing. While glucose and gluconate are catabolized directly through activities within the EDEMP cycle (21), fructose is transported and phosphorylated by means of a dedicated sugar phosphotransferase (PTS) system (27), and ribose feeds the non-oxidative PP route (28). Citrate was chosen as an entirely gluconeogenic substrate (29), whereas glycerol enters at an intermediate point in the catabolic network of *P. putida*, triggering a mixed gluconeogenic and glycolytic processing regime (30). Glycerol enters into the primary metabolism *via* its conversion to glycerol-3P **(Fig. 2A)** and then dihydroxyacetone phosphate and glyceraldehyde-3P (GA3P). From there, C3 units are metabolized either through the lower glycolysis [GA3P→pyruvate (Pyr)] or the gluconeogenic arm of the EMP pathway to generate intermediates with longer carbon backbones (GA3P→hexoses phosphate) (30, 31). Citrate enters directly into the tricarboxylic acid (TCA) cycle **(Fig. 2A)** and, besides direct oxidation, gluconeogenesis is used for biosynthesis of longer carbon chain metabolites (32).

**Figure 2.**
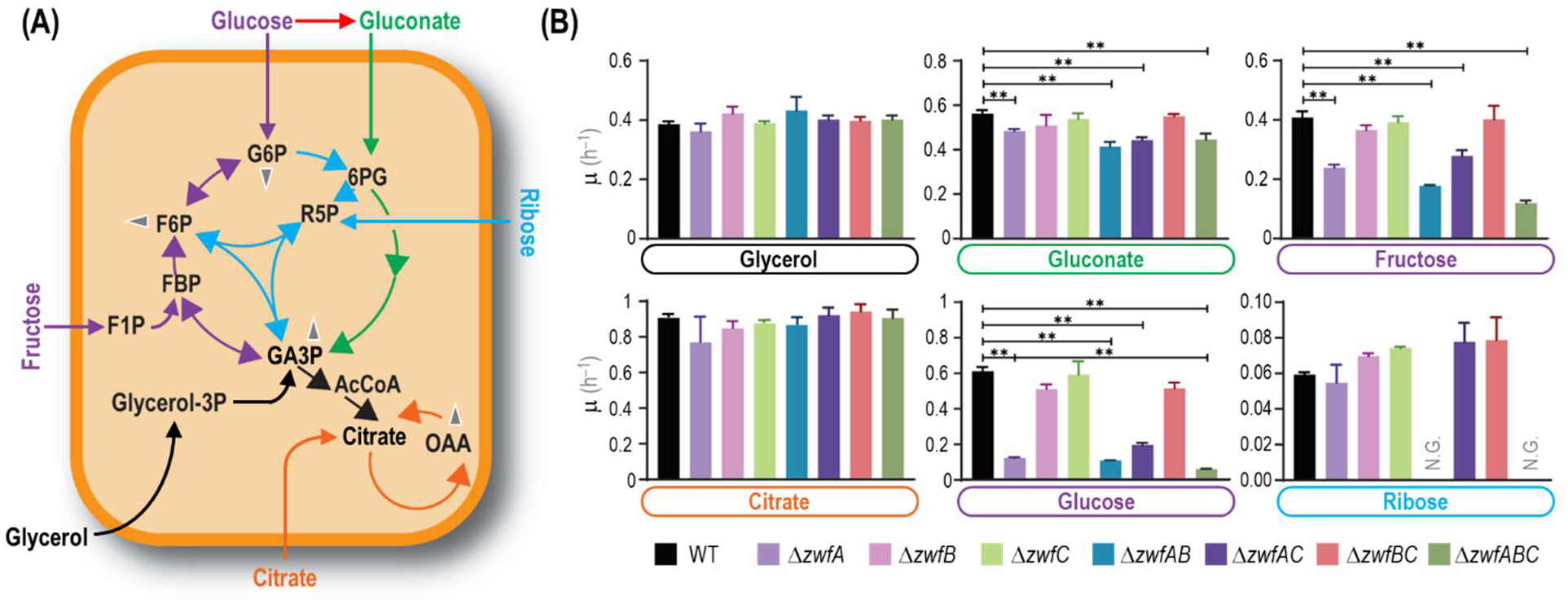
Analysis of growth patterns of different *zwf* mutants of *P. putida* KT2440. **(A)** Simplified scheme of the active metabolic blocks in the biochemical network of *P. putida* when cells are grown on the carbon sources indicated. Color codes and abbreviations are as indicated in the legend to **Fig. 1**. Glucose and fructose (C6) directly feed the (incomplete) EMP pathway, gluconate (C6) enters the ED route and ribose (C5) is processed through the PP pathway. Citrate (C6) acts as an entirely gluconeogenic substrate, whereas glycerol (C3) is processed *via* both the upper gluconeogenic domain of the network and downward catabolism. Grey arrowheads indicate further conversion of key metabolic intermediates (e.g. used as precursors for biomass). **(B)** Batch cultures of wild-type (WT) strain KT2440 and all combinations of *zwf* deletion mutants were grown in M9 minimal medium supplemented with the substrates indicated such they provide 120 mM total carbon. Carbon sources are identified with different colors according to the metabolic block they feed. Specific growth rates (μ) were determined during exponential growth, and bars represent the mean μ value ± standard deviation from three independent experiments per substrate. Double asterisks (**) identify statistically significant differences at the *p* < 0.01 level, assessed with the homoscedastic Student’s *t* test. N.G., no growth.

All strains exhibited essentially the same growth patterns in glycerol and citrate cultures **(Fig. 2B)**, with μ values similar to those reported previously (30). The apparent lack of any effect of *zwf* deletions on μ in glycerol cultures came as a surprise, since a major fraction of the carbon flux (73-104% of the glycerol uptake) is known to be routed through G6PDH (33)—and also because the expression of the *zwfA*-*pgl*-*eda* operon is upregulated during growth of strain KT2440 on glycerol (24). These results indicate that *P. putida* has the ability to re-route the carbon flow during the growth on glycerol, sustaining the supply of anabolic precursors produced in the PP pathway even when the G6PDH reaction is absent. On the other hand, D’Arrigo et al. (32) noted that in citrate-grown *P. putida*, fructose-6-phosphate seems to enter the PP route directly—rather than being converted into G6P and subsequently entering the oxidative phase of the PP pathway. Consistent with such observation, the *zwf* mutants did not show differences in growth rates when growing on citrate **(Fig. 2B)**.

In contrast to the results above, μ values differed across the strains tested when grown on glycolytic carbon sources. Mutants lacking *zwfA* displayed significant growth defects in gluconate cultures **(Fig. 2B)**, a substrate that is oxidized in the periplasm before entering the ED pathway downstream of G6PDH. However, no significant reduction in μ was observed for mutants lacking *zwfB*, *zwfC* or both, indicating that the isoenzymes encoded by such genes do not display a significant role in gluconate metabolism. Gluconate assimilation highly induces the expression of genes in the HexR regulon (24) and, as a consequence, *zwfA* expression is upregulated. Indeed, our results suggest that the activity of G6PDH-A alone can support all the required G6PDH flux under these cultivation conditions. Since no G6PDH activity is needed during the first steps of gluconate assimilation, the effect of deleting *zwfA* in gluconate cultures should be related to the insufficient G6PDH activity for flux cycling through the EDEMP pathway provided by G6PDH-B and G6PDH-C.

While the differences in gluconate-dependent growth of *zwfA* mutants were relatively minor, the growth of some knock-out strains was severely impaired when cultivated on glucose or fructose. Besides a strong reduction in growth rates (e.g. μ^Δ*zwfA*^ was 84% lower than μ^KT2440^, **Fig. 2B**), an extended lag phase was also observed in some cases (data not shown). Considering the high flux through the G6PDH reaction, the high level of expression of *zwfA* observed in *P. putida* when growing on glucose (24), it was not surprising to observe very low μ values in mutants lacking *zwfA*. According to metabolic fluxes estimations, most of glucose is firstly oxidized to gluconate (21, 23). Therefore, differences observed between growth patterns of *zwf* mutants on glucose (strong phenotype) or gluconate (mild phenotype) deserve special attention. A likely explanation stems from the steady state assumed for both flux balance analysis and ^13^C-based metabolic flux analysis, which differs from what actually happens during batchwise growth of *P. putida* without carbon limitation. Dynamic periplasmic conversion of glucose into gluconate and 2-ketogluconate affects flux distributions in the cytoplasm. As such, differences in μ values in gluconate and glucose cultures are connected to the complex dynamics of glucose assimilation [e.g. delayed gluconate uptake (34)]. This occurrence could explain longer lag phases of glucose cultures, as well as the larger impact of deleting *zwf* when the mutants are grown on glucose.

Fructose is phosphorylated to fructose-1-P and subsequently to fructose-1,6-bisphosphate (FBP) (35) **(Fig. 2A)**. The single *zwfA* knockout displayed a significant decrease in μ **(Fig. 2B)** and, even though the elimination of *zwfB* or *zwfC* did not result in major growth defects, the double Δ*zwfAB* mutant and the triple Δ*zwfABC* mutant had synergistic effects of individual deletions—with μ^Δ*zwfABC*^ being 70% lower than μ^KT2440^. As can be deduced from the observations of the cultures using citrate or glycerol as the sole carbon sources, G6PDH activity can be fully substituted to feed the PP pathway. A possible explanation for the large growth difference of the same mutants on fructose as compared to glycerol is metabolic imbalance—as *P. putida* utilizes FBP aldolase in the gluconeogenic direction (36, 37). Therefore, *P. putida* prioritizes the ED pathway for metabolization of fructose over direct formation of GA3P and DHAP (38). As the equilibrium of FBP aldolase lies heavily to the side of FBP (39), high FBP concentrations are therefore needed to support a glycolytic flux through the EMP pathway (40).

Finally, no growth of the double Δ*zwfAB* deletion mutant was observed on ribose **(Fig. 2B)**, indicating that the catabolism of this pentose substrate also relies on the EDEMP cycle and that the G6PDH-C has a near-zero activity. Because growth on ribose of wild-type *P. putida* was slow (μ ~ 0.06 h^−1^), we confirmed the mutual essentiality of G6PDH-A and G6PDH-B by growing the Δ*zwfAB* strain on M9 minimal medium agar plates containing ribose for 180 h **(Fig. S2)**. As observed in the liquid medium experiments, no growth deficiency was observed for single knock-outs—suggesting that the expression of either *zwfA* or *zwfB* ensures sufficient G6PDH activity to support the flux required for growth. This observation is in line with the expected catabolism of ribose, where two F6P molecules and one GA3P molecule are produced per three molecules of sugar. F6P cannot be metabolized further than G6P, therefore F6P, G6P and FBP may accumulate in this metabolic branch and will prevent further growth. In other *Pseudomonas* species, the build-up of F6P seems to be relieved by alginate accumulation (41), but such mechanism is apparently not relevant in strain KT2440. In any case, and since the analysis of single and combined *zwf* deletions indicated a differential role in the catabolism of glycolytic and gluconeogenic carbon sources, the next step was assessing the kinetic properties of each isoform.

### Kinetic properties of individual G6PDH isoforms of P. putida KT2440 indicate different physiological roles

Although most G6PDH are described as NADP^+^-dependent enzymes, previous reports suggested that dual or even NAD^+^-preferring homologues exist in several bacterial species, including *Pseudomonas* and close relatives (17–19, 42–44). The kinetic properties of the G6PDH-B and G6PDH-C isoforms present in strain KT2440 have not been studied so far, and whether G6P oxidation catalyzed by these variants would funnel electrons into the NADH or NADPH pools remains unclear. Because each redox cofactor has a distinct metabolic function, we reasoned that assessing the relative production of nucleotides by different G6PDHs *in vitro* would help understanding the presence of isoforms and their respective roles in *P. putida*. To this end, the genes encoding G6PDH-B and G6PDH-C were cloned and expressed in *E. coli*, and the enzymes were purified to homogeneity by Ni^2+^-based affinity chromatography. The kinetic properties of these purified G6PDH variants were then studied using either NAD^+^ or NADP^+^ as the redox cofactors. The key biochemical parameters for G6PDH-A, G6PDH-B and G6PDH-C, including the turnover constant (*k_cat_*) and the Michaelis constant (*K_M_*) values for each variant, are reported in **Table 1**.

**Table 1.**
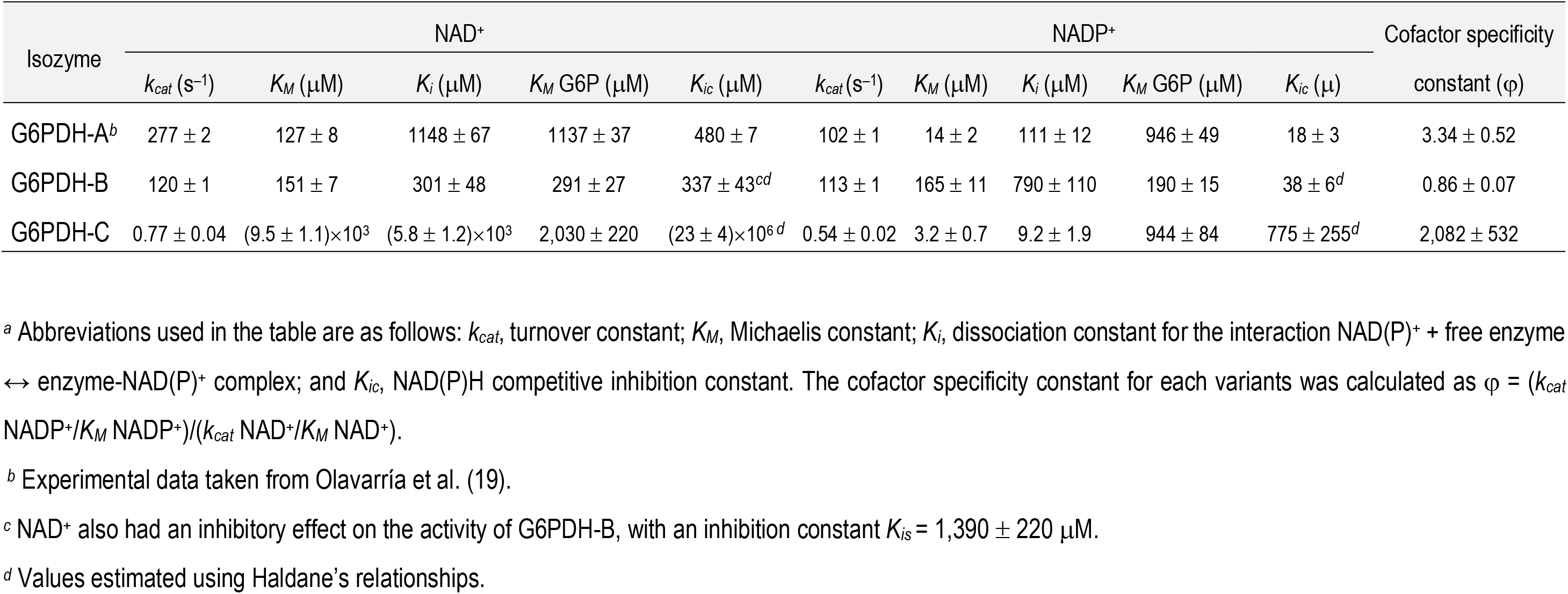
Kinetic parameters of the three G6PDH isoforms in *P. putida* KT2440^*a*^.

| Isozyme | NAD<sup>+</sup> |  |  |  |  | NADP<sup>+</sup> |  |  |  |  | Cofactor specificity |
| --- | --- | --- | --- | --- | --- | --- | --- | --- | --- | --- | --- |
| | $k_{cat}$ (s<sup>-1</sup>) | $K_M$ (μM) | $K_i$ (μM) | $K_M$ G6P (μM) | $K_{ic}$ (μM) | $k_{cat}$ (s<sup>-1</sup>) | $K_M$ (μM) | $K_i$ (μM) | $K_M$ G6P (μM) | $K_{ic}$ (μ) | constant (φ) |
| G6PDH-A<sup>b</sup> | 277 ± 2 | 127 ± 8 | 1148 ± 67 | 1137 ± 37 | 480 ± 7 | 102 ± 1 | 14 ± 2 | 111 ± 12 | 946 ± 49 | 18 ± 3 | 3.34 ± 0.52 |
| G6PDH-B | 120 ± 1 | 151 ± 7 | 301 ± 48 | 291 ± 27 | 337 ± 43<sup>cd</sup> | 113 ± 1 | 165 ± 11 | 790 ± 110 | 190 ± 15 | 38 ± 6<sup>d</sup> | 0.86 ± 0.07 |
| G6PDH-C | 0.77 ± 0.04 | (9.5 ± 1.1) × 10<sup>3</sup> | (5.8 ± 1.2) × 10<sup>3</sup> | 2,030 ± 220 | (23 ± 4) × 10<sup>6</sup> <sup>d</sup> | 0.54 ± 0.02 | 3.2 ± 0.7 | 9.2 ± 1.9 | 944 ± 84 | 775 ± 255<sup>d</sup> | 2,082 ± 532 |

<sup>a</sup> Abbreviations used in the table are as follows: $k_{cat}$ , turnover constant; $K_M$ , Michaelis constant; $K_i$ , dissociation constant for the interaction NAD(P)<sup>+</sup> + free enzyme ↔ enzyme-NAD(P)<sup>+</sup> complex; and $K_{ic}$ , NAD(P)H competitive inhibition constant. The cofactor specificity constant for each variants was calculated as $\phi = (k_{cat} \text{ NADP}^+ / K_M \text{ NADP}^+) / (k_{cat} \text{ NAD}^+ / K_M \text{ NAD}^+)$ .
<sup>b</sup> Experimental data taken from Olavarria et al. (19).
<sup>c</sup> NAD<sup>+</sup> also had an inhibitory effect on the activity of G6PDH-B, with an inhibition constant $K_{is} = 1,390 \pm 220$ μM.
<sup>d</sup> Values estimated using Haldane's relationships.

The rapid-equilibrium random ordered mechanism was the best model to explain the results observed in all cases, similarly to previous findings reported for the G6PDH-A isoform of strain KT2440 (19). Importantly, the main differences between G6PDH-A and G6PDH-B were detected in the *K_M_* values for NADP^+^ and G6P, which indicate that the response of these enzymes to changes in the cytoplasmic concentrations of these metabolites could be different **(Table 1)**. The cofactor specificity constants [φ, defined as (*k_cat_*^NADP+^/*K_M_*^NADP+^)/(*k_cat_*^NAD+^/*K_M_*^NAD+^)] were clearly different with respect to the φ of well-characterized NADP^+^-preferring homologues. G6PDH-C had a very different set of kinetic parameters when compared to the other two isozymes: this isozyme exhibited the highest *K_M_* (>10^3^ μM) for NAD^+^ and the lowest *K_M_* for NADP^+^ (< 10 μM) among the three G6PDHs. Therefore, G6PDH-C shows a very high specificity for NADP^+^ [as a comparison, the G6PDH from *E. coli*, widely regarded as an NADP^+^-preferring dehydrogenase (45), is reported to have a φ = 410, while φ^G6PDH-C^ is 5-fold higher than that value). However, it exhibited a very slow turnover (*k_cat_* < 1 s^−1^), both in the presence of NAD^+^ and NADP^+^. Such a low turnover constant is consistent with the negligible effect of deleting *zwfC* on μ under all cultivations conditions tested **(Fig. 2B)**.

Considering the results above and especially the differences in φ values among the dehydrogenase variants, we addressed the question of the cofactor specificity under *in vivo*-like conditions. Although φ provides a clear indication of cofactor preference under saturating conditions, physiologically-relevant circumstances are dynamic and often different to the setup adopted for *in vitro* kinetic assays. One of the most important differences *in vivo* is the simultaneous presence of NAD^+^, NADP^+^, NADH and NADPH— besides varying concentrations of G6P and other potential effectors not added to the *in vitro* assays. Thus, we applied a simplified kinetic model to evaluate the relative production of NADPH and NADH by the G6PDH isozymes under study under physiologically-relevant conditions, i.e. considering variable G6P concentrations and NAD^+^/NADH and NADP^+^/NADPH redox ratios consistent with the thermodynamic constrains enabling the operation of metabolism **(Fig. 3)**. Significant differences between the G6PDH isoforms were observed when the relative NADH-to-NADPH output was simulated as a function of the redox ratios at two G6P concentrations spanning one order of magnitude [i.e. 100 μM and 1 mM, compatible with values reported for glucose-grown strain KT2440 (21, 22)]. While the activity of G6PDH-A yields essentially the same amounts of NADH and NADPH regardless of the G6P concentration (NADH produced per NADPH close to 1) **(Fig. 3A)**, G6PDH-B generated mostly NADH and the relative production of NADH over NADPH showed a higher dependence on theG6P concentration **(Fig. 3B)**. The high sensitivity to substrate availability comes from the fact that G6PDH-B has a *K_M_* for G6P 5-fold lower than does G6PDH-A, either with NAD^+^ or NADP^+^ as cofactor **(Table 1)**. G6PDH-C, in turn, has the lowest NADH-to-NADPH output across the experimental conditions simulated herein **(Fig. 3C)**, consistent with its high φ. The notion that NADP^+^ is the preferred cofactor of the archetypal G6PDH^*E. coli*^ becomes evident when its NADH-to-NADPH output profile is compared to G6PDH-A and G6PDH-B **(Fig. 3D)**. Moreover, the relative NADH over NADPH production of G6PDH^*E. coli*^ has a more pronounced dependence on the G6P concentration. These biochemical features indicate different roles in the cellular redox balance for the different G6PDH isozymes. Yet the regulatory pattern of the cognate genes (and, in particular, *zwfC*) remains unclear—and we set out to investigate this aspect as disclosed in the next section.

**Figure 3.**
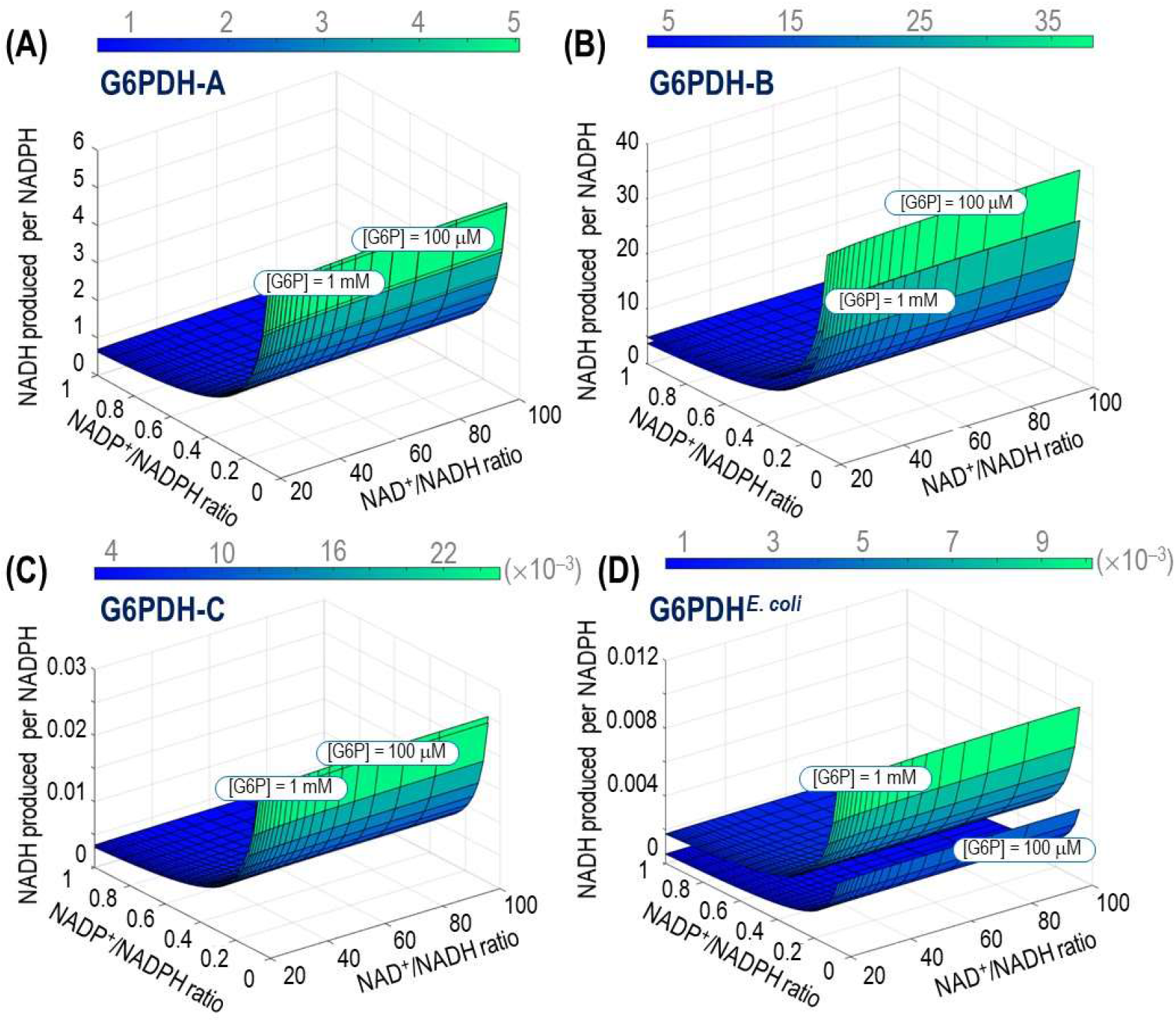
Relative NADH-to-NADPH output of the G6PDH isozymes of *P. putida* KT2440. The relative production of the two reduced cofactors was simulated across physiologically-relevant NAD^+^/NADH and NADP^+^/NADPH ratios and two glucose-6-phosphate (G6P) concentrations for **(A)** G6PDH-A, **(B)** G6PDH-B and **(C)** G6PDH-C. The relative NADH-to-NADPH production was also simulated for the single G6PDH enzyme of *E. coli*, included here for the sake of comparison **(D)**.

### The zwfC gene from P. putida KT2440 is poorly transcribed across different growth conditions

According to the experiments described previously, the deletion of *zwfC* has no significant effect on the growth patterns in any of the carbon sources tested **(Fig. 2B)**. Therefore, we wondered whether *zwfC* encodes a functional G6PDH enzyme—especially considering that a purified G6PDH-C showed the highest specificity for NADP^+^ *in vitro* **(Table 1)**. According to the genome annotation, *zwfC* encodes a non-truncated G6PDH-like protein. Additionally, an alignment of G6PDH-C with the G6PDH enzyme from *Leuconostoc mesenteroides* revealed that both the cofactor binding motif [G_12_-(G/A)-X-GDL-(A/V)-(K/L) at the *N*-terminus, with X representing any amino acid] and other residues key for the interaction with substrates (i.e. R176IDHYLGKE, E146KP, Y415 and H240) are conserved. However, its expression and/or activity could be regulated by transcriptional, translational or post-translational mechanisms that could render the gene or the polypeptide silent. To explore this possibility at the transcriptional level of regulation, we first searched for ribosomal binding sites (RBS) upstream of the open reading frame of *zwfC* by using the RBS calculator tool (46). Several in-frame *START* codons, with a predicted medium-to-strong translation efficiency, were found around the annotated start of the open reading frame according to this analysis. Secondly, we looked for potential promoter regions upstream of the G6PDH-C encoding sequence. Interestingly, the 5’-untranslated region of *zwfC* had a duplicated HexR binding motif, highly similar to the recognition motif found upstream of *zwfA* **(Fig. 4A)**. This motif follows the canonical consensus HexR-binding site **(Fig. 4B)**, which was originally identified by Daddaoua et al. (47) not only for *zwfA*, but also for *gapA* and *edd*. Furthermore, upstream of *zwfC* there is a divergent gene, *PP_5350* (867 bp), predicted to encode a transcriptional regulator of the RpiR family **(Fig. 4C)**. This RpiR regulator shares a high similarity with HexR (45% identity, *hexR* is 864 bp long) and the binding moieties for sugars (residues R53 and R56) as well as for DNA (residues S139, S183) are conserved (48). Interestingly, such genomic organization is not unique to *P. putida*, and we have identified orthologues of *rpiR* and *zwfC* adjacent to each other in many annotated genomes of *Pseudomonas* and closely related species (representative examples are shown in **Fig. S3**). Such architecture, where the gene encoding a putative transcriptional regulator is located close to the regulated gene(s), is widespread in nature.

**Figure 4.**
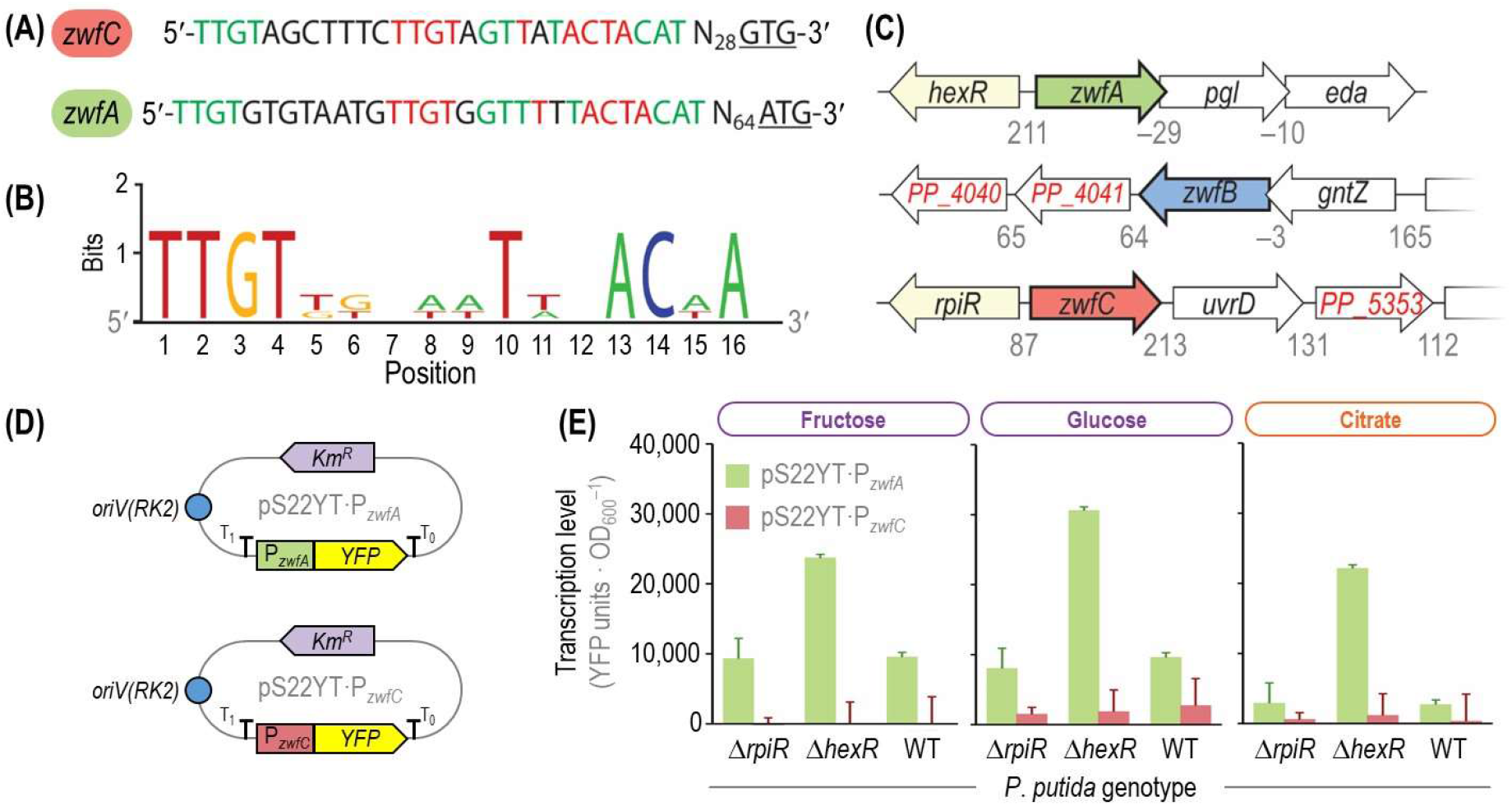
Genetic and genomic organization of *zwf* orthologues in *P. putida* KT2440 and analysis of transcriptional levels. **(A)** Upstream region of the *zwfA* and *zwfC* genes. Both regions contain a duplicated HexR-binding site (marked in red) and share sequences with high similarity (indicated in green) as annotated by Belda et al. (90). The start codons of *zwfA* and *zwfC* (*ATG* and *GTG*, respectively) are underlined, and N represents any nucleotide. **(B)** Consensus sequence of HexR-binding sites for *zwfA*, *gapA* (encodes glyceraldehyde-3-phosphate dehydrogenase) and *edd* (encodes 6-phosphogluconate dehydratase). Genomic organization of the three *zwf* orthologues of strain KT2440. The *zwfA* gene forms an operon together with *pgl* and *eda* (encoding 6-phosphogluconolactonase and 2-keto-3-deoxy-6-phosphogluconate aldolase, respectively), and its expression is controlled by the HexR regulator. The *zwfB* gene is co-transcribed with *gntZ* (encodes 6-phosphogluconate dehydrogenase), while *zwfC* is divergently transcribed with respect to *rpiR* (PP_5050, a transcriptional regulator). Numbers below the gene clusters indicate intergenic distances in base pairs, with negative values representing an overlap between the corresponding open reading frames. Elements in this diagram are not drawn to scale. Reporter plasmids constructed to assess *zwfA* and *zwfB* expression levels. The promoter region of the corresponding genes was cloned in front of the gene encoding the yellow fluorescent protein (YFP), and the resulting modules are transcriptionally insulated by means of the T_0_ and T_1_ terminators. These low-copy-number [*oriV*(RK2)] reporter plasmids are endowed with a kanamycin resistance (Km^R^) cassette. **(E)** Transcriptional analysis of the P_*zwfA*_ and P_*zwfC*_ promoters in different genetic backgrounds. Wild-type (WT) *P. putida* KT2440 and its Δ*hexR* or Δ*rpiR* mutant derivatives were transformed with the reporter plasmids described in **(D)**, carrying the promoter regions of *zwfA* and *zwfC*. Fluorescence values were normalized against those in the WT strain carrying vector pSEVA227Y (promoter-less YFP). Carbon sources are identified with different colors according to the metabolic block they feed (see **Fig. 2**). Bars represent mean values ± standard deviation from three independent experiments per substrate. OD_600_, optical density measured at 600 nm.

In order to investigate the potential role of RpiR on the expression of *zwfC*, we constructed a set of transcriptional reporter plasmids **(Fig. 4D)** derived from the low-copy-number pSEVA227Y vector (49). Plasmid pS22YT·P_*zwfC*_ contains the 278-bp-long 5’-UTR of *zwfC* (spanning the predicted promoter sequence) in front of the gene encoding for the yellow fluorescent protein (YFP). The upstream region of *zwfA* was also cloned in the same backbone, yielding plasmid pS22YT·P_*zwfA*_, where the promoter region of *zwfA* drives the expression of *YFP*. In addition, we created individual Δ*rpiR* and Δ*hexR* mutant derivatives of *P. putida* KT2440 **(Table S1)**. These deletion mutants, together with the wild-type strain, were transformed with the reporter plasmids or the empty pSEVA227Y vector and cultivated to measure the signal of the reporter protein across different growth conditions. No significant *YFP* expression from P_*zwfC*_ was detected in the wild-type strain or in the *ΔhexR* or *ΔrpiR* mutants when grown on fructose **(Fig. 4E)**. Detectable expression levels from the P_*zwfA*_ promoter were observed in all genetic backgrounds, with a strong (2.5-fold) increase in YFP fluorescence in the *ΔhexR* mutant compared to that of *P. putida* KT2440 **(Fig. 4E)**. A similar trend was also observed in glucose cultures. In this case, and even though expression from P_*zwfC*_ was marginally higher in all the tested strains, the differences in YFP levels in the wild-type strain and the mutants were not significant **(Fig. 4E)**. When the experiments were repeated in the presence of citrate, a gluconeogenic carbon source, almost no expression from P_*zwfC*_ was observed in any of the genetic backgrounds tested. The normalized YFP signal triggered by P_*zwfA*_ in citrate cultures, in contrast, was around half as strong as observed in the wild-type strain and the *ΔrpiR* mutant on fructose—and similarly high as in the *ΔhexR* mutant grown on fructose **(Fig. 4E)**. In agreement with these observations, low *zwfC* mRNA levels were detected by deep sequencing of the transcriptome of strain KT2440 grown on several carbon sources (50), while the highest *zwfA* transcription levels were observed on glucose **(Fig. S4)**. Taken together, these results indicate that the levels of *zwfC* expression are extremely low under either glycolytic or gluconeogenic regimes, and that the regulatory role of RpiR (if any) does not seem to be relevant in the experimental conditions tested. The next addressed question was how the G6PDH activity in the different zwf mutants is.

### G6PDH-A provides the bulk of the glucose-6-phosphate dehydrogenase activity in glucose-grown *P. putida* KT2440

The physiological data **(Fig. 2)**, the *in vitro* biochemical characterization of purified enzymes **(Table 1)** and the transcriptional pattern **(Fig. 4)** strongly suggest that G6PDH-C does not carry a significant flux, which prompts the question of which isozyme(s) actually provide the G6PDH activity in the strain KT2440. To this end, the G6PDH activities of all the constructed *zwf* mutant strains were assayed in cell-free extracts obtained from glucose and citrate cultures **(Fig. 5)**. Moreover, we tested enzyme activities using either NAD^+^ or NADP^+^ as the cofactor added to the reaction mixture. As expected, the overall G6PDH activity was consistently higher (around one order of magnitude) in cells grown under a glycolytic regime than in gluconeogenic conditions. The specific activity was almost lost in mutants deficient in G6PDH-A when grown on glucose, indicating that this isozyme supplies by far the most activity under glycolytic conditions. The specific G6PDH activities in the wild-type strain were similar to previous reports (21, 22), and the observed activity of the Δ*zwfABC* triple mutant in cell-free extracts was very low, indicating that only low promiscutive G6PDH of other proteins is present **(Fig. 5A)**. No activity above Δ*zwfABC* triple mutant levels could be detected in the Δ*zwfAB* double mutant, indicating, again, that G6PDH-C does not supply a significant G6PDH activity under the conditions tested. Interestingly, in glucose-grown cells, the G6PDH activity was essentially the same regardless of the cofactor used in the reactions, similar to the simulations shown in **Fig. 3A**, indicating that G6PDH-A is the predominant isozyme under such conditions. The bulk G6PDH activity in citrate-grown *P. putida* was in general lower that under glycolytic conditions **(Fig. 5B)**. The overall pattern was similar as indicated before, and G6PDH-A seems to carry most of the activity, but higher activity was observed with NADP^+^ as cofactor. These observations led us to investigate if the presence and cofactor preference of multiple G6PDHisoforms could be a redox adjusting mechanism in *Pseudomonas*.

**Figure 5.**
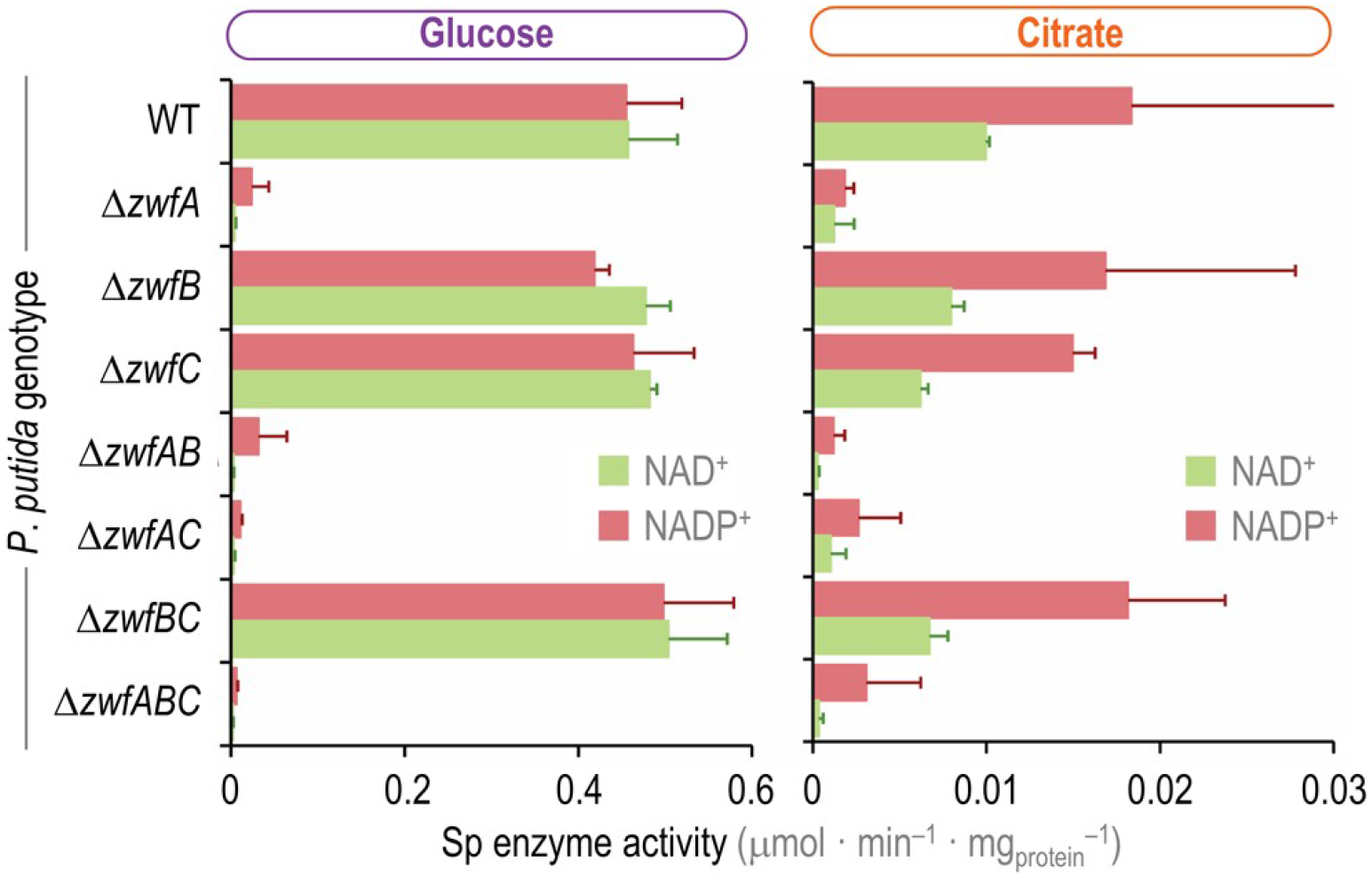
*In vitro* determination of glucose-6-phosphate dehydrogenase activity in *P. putida* KT2440 and the different *zwf* knock-outs. The specific (Sp) G6PDH activity was determined *in vitro* in cell-free extracts obtained from glucose or citrate cultures, assayed in the presence of either NAD^+^ or NADP^+^ as indicated. Carbon sources are identified with colors according to the metabolic block they feed (see **Fig. 2**). Data represent the mean value of the specific enzymatic activity ± standard deviation of triplicate measurements from at least three independent experiments.

### Genetically-encoded cofactor specificity of G6PDH isozymes in *P. putida* KT2440

An alignment of the amino acid sequences encoded by *zwfA*, *zwfB* and *zwfC* **(Fig. S5)** shows a relatively high degree of similarity: 54% between G6PDH-B and G6PDH-C, and 60% between G6PDH-A and G6PDH-C. However, the information gathered so far points to distinct kinetic properties, and a predominant role of G6PDH-A under the tested conditions. We therefor searched for signatures in the dehydrogenase polypeptides that could point to differences in cofactor specificity. One elegant example in this direction is the analysis of the (unique) G6PDH enzyme of the parasitic euglenoid *Trypanosoma cruzi* (51). Mercaldi et al. (52) reported that this G6PDH displays a key amino acid residue in the β2-α2 domain, R72, which specifically interacts with the 2′-phosphate group of NADPH **(Fig. 6A)**. Building on this rationale, this amino acid residue (arginine) was found to be determinant for the usage of NADP^+^ by the G6PDHs from several organisms, including *E. coli* (53) and the lactic acid bacterium *L. mesenteroides* (54, 55). Furthermore, the G6PDH isozymes of *Komagataeibacter hansenii*, *Burkholderia cepacia* and *Pseudomonas fluorescence* (biotype E), which are the most NAD^+^-specific G6DPH characterized so far, lack this specific arginine residue (16, 18, 44, 53, 56). Thus, we reasoned that if a given G6PDH contains an arginine residue at the position, predicted to interact with the 2′-phosphate moiety of NADP^+^, it can be considered either NADP^+^ or dual cofactor-specific, whereas other amino acids in this position would lower the NADP^+^-affinity of the enzyme.

**Figure 6.**
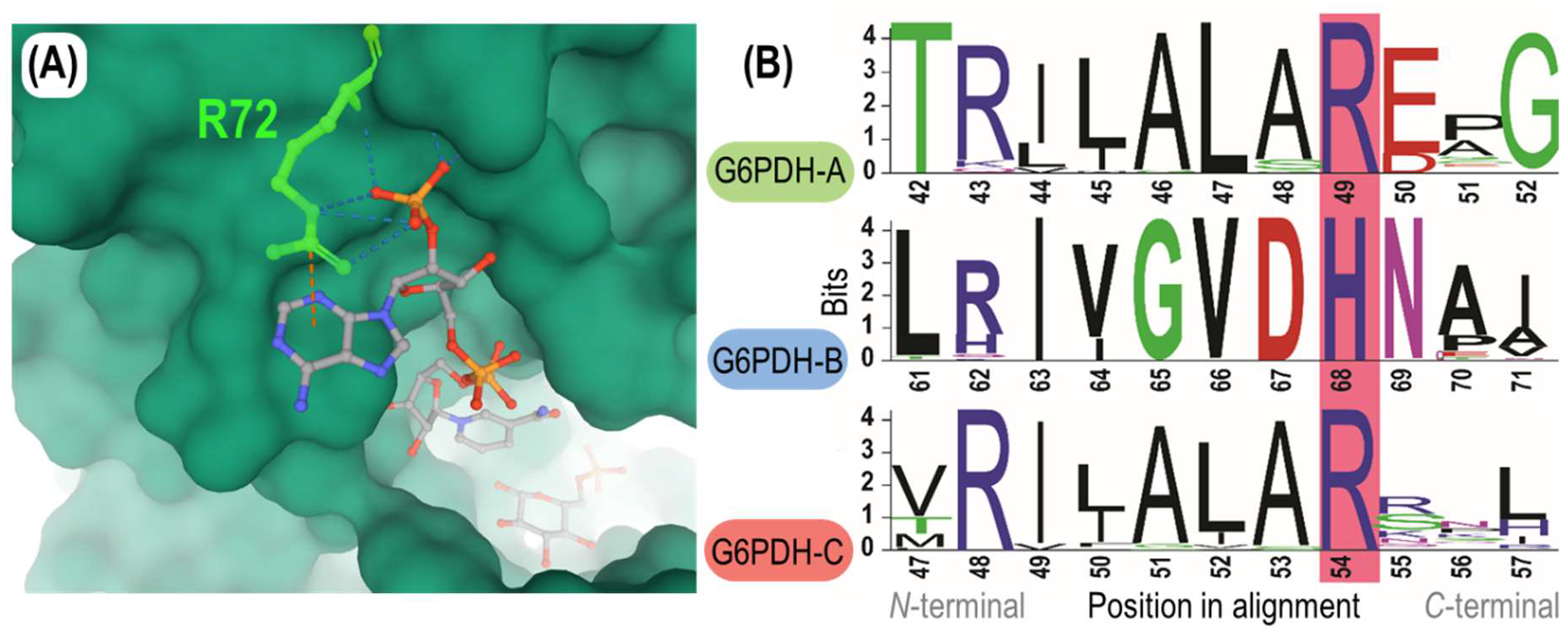
The β2-α2 domain region of the G6PDH isozymes of *P. putida* KT2440. **(A)** 3D model of the cofactor binding pocket of the G6PDH enzyme of *Trypanosoma cruzi*. R72, the cofactor discriminating residue, located in the β2-α2 loop of the protein and corresponding to R49 in G6PDH-A of *P. putida* KT2440, is highlighted in green. R72 interacts with the 2′-phosphate group of NADP^+^ through four hydrogen bonds, as indicated with broken lines in the diagram. Crystal structure adapted from Mercaldi et al. (52). **(B)** Alignment of the amino acid sequence of the three G6PDH forms of *P. putida* KT2440. G6PDH-A, G6PDH-B and G6PDH-C belong to family I, II and III, respectively, of glucose-6-phosphate dehydrogenases. Orthologue entries from OrthoDB (57) were aligned for each family (*N* = 101, 66 and 50, respectively), and amino acid residues defining cofactor specificity (G6PDH-A^R49^, G6PDH-B^H68^ and G6PDH-C^R54^) are highlighted in pink.

Following this reasoning, three distinct groups—each highly conserved around the β2-α2 loop of the enzyme—were identified when all the known G6PDH orthologues across the entire *Pseudomonas* genus available in orthoDB (57) were analyzed. The G6PDH isoforms of *P. putida* KT2440 fall each in one of these groups **(Fig. 6B)**: two of these groups contain the residue arginine in the β2-α2 loop (*i.e.* G6PDH-A^R49^ and G6PDH-C^R54^), while one family contains a conserved histidine in the cofactor-discriminating position (i.e. G6PDH-B^H68^). These results are in agreement with the *in vitro* characterization of purified enzymes **(Table 1)** and the φ values reported for each variant, as the presence of an arginine residue at the cofactor-discriminating position of G6PDH-A and G6PDH-C correlates with increased affinity towards NADP^+^. G6PDH-B, in contrast, displays a histidine in this loop, which matches the lack of specificity of this variant towards the redox cofactors NADP^+^ (φ^G6PDH-B^ ~ 0.9, **Table 1**). Taken together, these observations indicate that *P. putida* KT2440 harbors two main G6PDH functions relevant for *in vivo* conditions, G6PDH-A and G6PDH-B, and that these variants significantly differ in the cofactor specificity, probably enabling metabolic flexibility depending on the redox demand—which leads to the question of how widespread this correlation is across different bacterial species.

### The number and cofactor specificity of glucose-6-phosphate dehydrogenase isozymes correlate with the metabolic lifestyle across bacterial species

After identifying the presence of key residues defining NAD(P)^+^-acceptance in the G6PDH isozymes of strain KT2440, we were interested to explore if duplication of G6PDH-encoding genes is a widespread phenomenon in the bacterial domain and whether it has a connection with cofactor specificity. To this end, we searched all entries annotated as G6PDH in the OrthoDB (57), and clustered them according to phylogenetic relationships between species **(Fig. 7A)**. First, we observed that 88% of all analyzed *Pseudomonas* species harbor two or three G6PDH isozymes. Furthermore, we classified the corresponding isozymes as NADP^+^- or NAD^+^-dependent according to the presence or absence of the key arginine residue in the β2-α2 loop, respectively. While only 10% of the *Pseudomonas* species with a single *zwf* gene encode a NAD^+^-specific isoform, 65% of *Pseudomonas* with multiple *zwf* orthologues contain at least one NAD^+^-specific isozyme. Therefore, harboring multiple G6PDH isoforms with different cofactor specificities is an evolutionary trait common to the *Pseudomonas* genus. The analysis was subsequently broadened to the entire bacterial kingdom. Eukaryotes were excluded from this classification, as gene isoforms are known to be differentially expressed depending on the cell type or tissue (58). Archaea were likewise neglected, since archaeal G6PDH enzymes are not related to bacterial counterparts (59), which would make the cofactor specificity impossible to assign.

**Figure 7.**
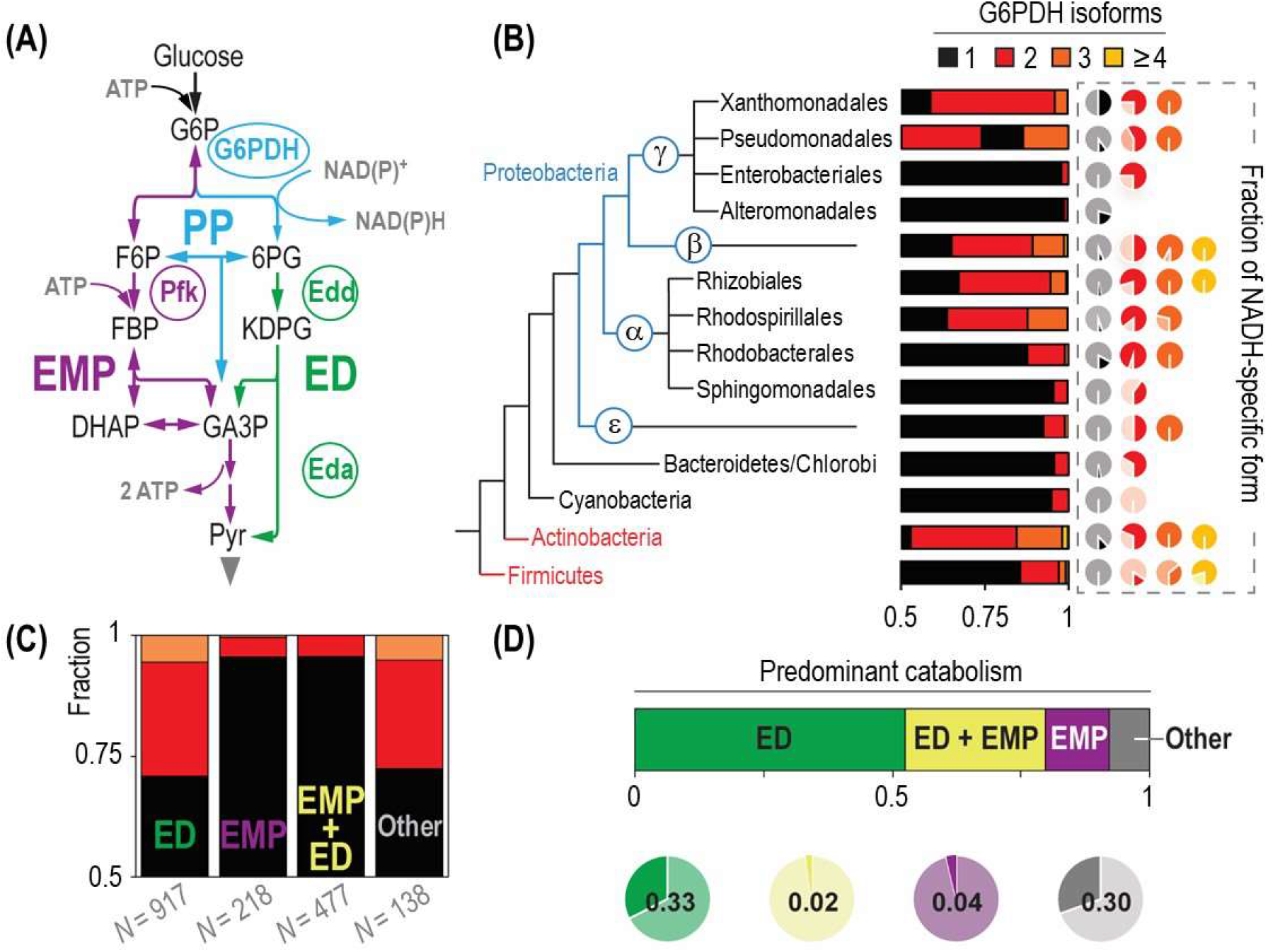
Cofactor specificity of G6PDH isoforms correlates with the metabolic lifestyle of the host. **(A)** Phylogenetic tree reconstruction of bacteria harboring different G6PDH variants. The fraction of microbial species in each order that encode one, two, three and four or more G6PDH isozymes is indicated together with the cofactor specificity of each of these fractions. NAD^+^ specificity was determined by the absence of the key arginine residue at the position interacting with the 2′-phosphate group of NADP^+^ (see **Fig. 6B**), and the fraction is indicated by a dark hue (a light shade is used for the NADP^+^-dependent variants). Information on *zwf* orthologues included in the analysis was gathered from OrthoDB (57). **(B)** Schematic comparison of the Entner-Doudoroff (ED) and Embden-Meyerhof-Parnas (EMP) glycolytic pathways. The EMP pathway converts glucose-6-phosphate (G6P) into fructose-6-phosphate (F6P), followed by a phosphorylation catalyzed by 6-phosphofructo-1-kinase (Pfk) that yields fructose-1,6-bisphosphate (FBP). Two molecules of glyceraldehyde-3-phosphate (GA3P) are finally formed through an aldolase and isomerization reaction. In the ED pathway, in contrast, G6P is oxidized to 6-phosphogluconate (6PG) through G6PDH (yielding a reducing equivalent) and 6-phosphogluconolactonase. After 6PG oxidation to 2-keto-3-deoxy-6-phosphogluconate (KDPG) by 6-phosphogluconate dehydratase (Edd), one GA3P molecule and one pyruvate (Pyr) molecule are formed by KDPG aldolase (Eda). Note that the EMP pathway produces twice the ATP than the ED pathway. Color coding of different metabolic blocks and other abbreviations used herein are as indicated in the legend to **Fig. 1**. **(C)** Correlation between glycolytic pathways of the species analyzed and the number of *zwf* genes they harbor. The fraction of species relying solely on the ED pathway, both the ED and EMP pathways, or solely on the EMP pathway was established by assessing the presence of unique genes for each metabolic block. The cases where no genes of the ED or the EMP pathway could be found were categorized as *others*. The color code identifying the number of *zwf* orthologues is the same as for **(B)**. **(D)** Correlation between the cofactor preference of G6PDH isozymes and the catabolic lifestyle of the hosts. The fraction of NAD^+^-specific G6PDH forms is plotted according to the predominant glycolytic route (or their combination). The Actinobacteria and Firmicutes phyla, highlighted in red in the phylogenetic tree **(B)**, are excluded from the analysis shown in **(C)** and **(D)**.

Many bacterial species were found to harbor multiple *zwf* genes **(Fig. 7A)**, but these orthologues are not homogenously distributed throughout all bacterial orders. While some orders (especially Proteobacteria) were observed to harbor a single G6PDH isoform almost exclusively (e.g. *Enterobacteriales* and *Alteromonadales*), almost 50% of species in other orders, e.g. *Xanthomonadales* or Actinobacteria, were found to encode multiple G6PDH variants. Within the proteobacterial domain, only α- (*Rhizobiales*) and β-proteobacteria encoded ≥ 4 G6PDH isozymes in their genomes (Actinobacteria and Firmicutes also shared this trait). In order to investigate the cofactor specificity of all these G6PDHs, the amino acid sequences of all enzymes was aligned and analyzed according to the presence of the key residue discriminating NADP^+^. The amount of organisms harboring a NAD^+^-specific G6PDH is very low, if the corresponding species has a single copy of *zwf*, but this fraction increases drastically in bacteria with multiple G6PDH isoforms **(Fig. 7A)**. The increase in the fraction of NAD^+^- to NADP^+^-preferring dehydrogenases is much higher than what would be expected just by chance. The general trend is that bacterial species seem to have a NAD^+^-specific isoform if they contain multiple G6PDH variants—while single G6PDH isoforms appear to be NADP^+^-dependent or dual cofactor-specific. Given the heterogeneous distribution across bacterial orders, we wondered whether the multiplicity of G6PDH isoforms and their redox cofactor preference might correlate with the metabolic lifestyle of these species.

To investigate this possibility, all bacterial species were categorized according to the presence of signature glycolytic routes. According to the simplified classification that we implemented to this end, organisms harboring 6-phosphofructo-1-kinase (Pfk) were considered capable of running the EMP pathway, while 6-phosphogluconate dehydratase (Edd) or 2-keto-3-deoxy-6-phosphogluconate (KDPG) aldolase (Eda) was chosen as markers of the ED pathway **(Fig. 7B)**. If none of these genes was found in a given species, the glycolytic strategy of the organism was labeled as “other”—a category that includes, among others, phosphoketolase-dependent and entirely fermentative lifestyles. We observed that the prevalence of multiple G6PDH variants strongly correlates with the use of the ED pathway as the sole glycolytic route in all orders **(Fig. 7C)**, with the exception of the Actinobacteria and Firmicutes phyla. If these orders are excluded from the analysis, we observed that 29% of microbes relying on the ED pathway harbor multiple G6PDH isoforms—in contrast to just 4.5% and 4.4% for EMP- and EMP and ED-utilizing organisms, respectively. Also, 28% of organisms lacking key enzymes of both EMP and ED pathways display multiple G6PDH isoforms. Therefore, the emerging picture is that bacteria that do not possess the capacity of running the EMP pathway show a higher fraction of NAD^+^-specific G6PDH variant. This is in agreement with previous findings (16). Furthermore, we confirmed that all glycolytic routes are well represented in this analysis **(Fig. 7D)**, considering that the query for bacteria containing G6PDH enzyme(s) could potentially create a bias towards microbes using the ED route. A similar trend was observed for the cofactor specificity of the G6PDH isoforms **(Fig. 7D)**: 33% and 30% of organisms using solely the ED pathway or neither the ED nor EMP routes were observed to harbor a non-NADP^+^– specific G6PDH variant (i.e. NAD^+^-preferring), respectively. This observation contrasts with the percentage of bacteria relying either solely on the EMP pathway or the EMP and ED routes that carry a NAD^+^-specific G6PDH variant (4% and 2%, respectively).

A clear outlier in this analysis are the members of Actinobacteria **(Fig. 7A)**, and they deserve a separate discussion. The majority of organisms in the Actinobacteria phylum contain the necessary genes for both the EMP and the ED pathway. Therefore, a relatively low abundance of organisms with multiple isoenzymes and NAD^+^-dependent variants would have been expected according to the interpretation above. However, we observed that almost half of the species have multiple *zwf* genes and NADH-yielding isozymes are relatively abundant. Actinobacteria are not related to the other phyla analyzed in this study, and they might have evolved different metabolic strategies. Moreover, it appears that the EMP pathway is preferred over the ED and PP pathway for glucose catabolism in this phylum. *Streptomyces coelicolor* diverts around half the glycolytic flux through the EMP route and half through the ED pathway (60). Similarly, 62% and 52% of the glucose uptake flux is directed through the reaction catalyzed by the G6PDHs in *Corynebacterium glutamicum* and *Streptomyces lividans*, respectively (61, 62). *Nonomuraea* sp. ATCC39727 even uses the ED pathway predominantly, although the genes encoding for the enzymes of the EMP pathway are present in the genome (63). Thus, the high flux through G6PDH even when the EMP pathway is present explains the high abundance of NADH-yielding isoforms in this phylum.

## Discussion

The notion that enzymes became highly specialized to execute a single function is often misleading— many of them exhibit a wide spectrum of substrates leading to accuracy-rate tradeoffs, thereby affecting evolutionary trajectories (64). We examined this issue in the model environmental bacterium *P. putida*, and we found that the three G6PDH isozymes of strain KT2440 display different cofactor specificity. While G6PDH-A (the production of which is induced by glucose) uses NADP^+^ and NAD^+^ with similar proportions, G6PDH-B (which is constitutively expressed) prefers NAD^+^. According to the physiological characterization of mutants lacking *zwfA*, this variant displays the most prominent role in batch cultures with glucose as the carbon source, where substrate abundance in the medium is high and cytoplasmic G6P concentrations are typically elevated (21, 65). This situation helps explaining the higher *K_M_* of G6PDH-A for G6P as compared to the other two variants, and hints to an important contribution of G6PDH-B to catabolism during carbon limitation. The interplay between the activities afforded by G6PDH-A and G6PDH-B appears to balance NADPH production on different carbon sources, since only a small fraction of the carbon flux is directed through G6PDH during glucose-dependent growth—whereas almost the entire flux is funneled through this reaction when the cells are grown on fructose or ribose.

G6PDH-C, in contrast, has an extremely low turnover both *in vivo* and *in vitro* and its deletion had negligible effects on bacterial growth. Thus, the function of this isozyme remains elusive—and its role could even be structural or regulatory, rather than metabolic. Examples of this sort (where dehydrogenase enzymes display a non-metabolic function) include one of the GA3P dehydrogenase isozymes of the pathogen *Neisseria meningitidis*, which plays an important role in the adhesion of the bacterium to host cells (66). Another instance is the structural role of lactate dehydrogenase in the lenses of crocodiles and birds (67). While it is possible that G6PDH-C requires protein(s) and/or allosteric effector(s) absent in our enzymatic assays to be rendered fully functional, this is usually not the case for this type of dehydrogenase activity. Another possibility is that this enzyme prefers another sugar substrate different from G6P, as demonstrated for the sulfoquinovose catabolic pathway of *P. putida* SQ1 (68), or an alternative redox cofactor besides NAD(P)^+^. These scenarios notwithstanding, we hypothesize that the existence of G6PDH-A and G6PDH-B as the main metabolism-related G6PDH isozymes “freed” G6PDH-C to explore new functions. This phenomenon (also known as ‘moonlighting’) is a well-known feature of catabolic modules in environmental bacteria, which acquired the capability of processing alternative substrates *via* gene duplication and enzyme specialization, usually in connection to a local transcriptional factor (69, 70)—as observed for the gene encoding the RpiR-like regulator and *zwfC*. From a more general perspective, the so-called “patchwork” model theorizes that primitive enzymes were highly promiscuous to confer a larger degree of catalytic versatility when the pool of available biocatalysts was limited (71, 72). Besides the rich panoply of biochemical reactions, *P. putida* may bear underground metabolic pathways as a basis of its remarkable capacity to adapt to changing environments (73).

One way or the other, it seems that G6PDH-A and G6PDH-B carry most of the G6PDH activity in this species. The use of multiple such isoenzymes is prevalent in organisms lacking a functional EMP pathway and correlates with the use of NAD^+^-specific variants. We argue that this occurrence confers metabolic flexibility to the host without altering carbon fluxes. The dual cofactor specificity of G6PDH-B probably balances redox cofactor production in a narrow window just by demand, as the NADPH/NADH ratio associated to G6PDH activity is highly dependent on substrate (i.e. NADP^+^ and NAD^+^) availability (19). An increase in the demand of a specific cofactor leads to a higher concentration of its oxidized form, thereby shifting substrate availability and, subsequently, the cofactor output. The two active isozymes warrant a broader window for cofactor balancing to *P. putida*, and may provide a reserve flux capacity under oxidative stress. Both G6PDH-A and G6PDH-B are present in the cells simultaneously under all the experimental conditions investigated herein. G6PDH-A is inhibited by NADPH, with a K_i_ (NADPH) = 112 μM, lower than the intracellular NADPH concentration of 276 μM observed in strain KT2440 when grown under non-stressed conditions (74). A higher flux is needed to compensate for this inhibition and, if G6PDH-B supplies the extra activity, NADH would be produced. Under oxidative stress, the concentration of NADPH decreases and releases the inhibition on G6PDH-A, thus increasing the flux through this node and thereby producing NADPH. Furthermore, the different metabolic specialization of the two variants is also reflected by the lower *K_M_* of G6PDH-B for G6P than G6PDH-A (around 5-fold), thus favoring G6P processing through G6PDH-B in metabolic states with low substrate levels—e.g. growth on gluconeogenic carbon sources.

From a broader perspective, the correlation between the predominant glycolytic strategy of a given microorganism and the presence of G6PDH isozymes displaying different cofactor specificity seems to be largely dictated by the need of balancing the redox status on different carbon sources according to environmental conditions. As indicated above, bacteria relying on the EMP pathway for sugar catabolism can increase flux into the PP or the ED pathway through G6PDH whenever an extra supply of NADPH is needed. This is actually the case under oxidative stress conditions, as previously shown in *E. coli*, yeast and mammalian cells (10, 75, 76). Such a strategy is obviously not possible in microbes solely relying on the ED pathway, and the use of isozymes with different cofactor specificity becomes more prominent. Other redox-adjusting mechanisms include the use of peripheral (oxidative) pathways in the first steps of sugar processing, e.g. the gluconate/2-ketogluconate loop of *P. putida* **(Fig. 1)**, together with allosteric control of enzyme activities at key metabolic steps. Only 20% of the glucose is directly phosphorylated to G6P in *P. putida* (before being channeled through G6PDH), whereas 80% of the sugar is oxidized in the gluconate/2-ketogluconate loop and thus bypasses G6PDH (21). Accordingly, *P. putida* strongly expresses *zwfA*, encoding the NADPH-forming variant, under this condition—thereby adjusting the NADPH output by tuning the flux split between glucose phosphorylation and oxidation. Besides, the metabolism of glucose generates 24 reducing equivalents per molecule, while gluconate yields 22 reducing equivalents (77)—thus, the conversion of glucose to Pyr requires one additional oxidation step in comparison to gluconate. This adds a further layer of transcriptional control on the *zwf* orthologues of *P. putida*: expression of *zwfA* is enhanced in the presence of glucose to ensure this additional oxidation step, which is not required for the oxidation of gluconate. Given the absence of any overflow metabolism, this bacterium achieves redox balancing through the activity of alternative dehydrogenases rather than by producing fermentation products. These observations will also open up new strategies towards engineering metabolism of this species by harnessing the wealth of enzymatic activities typical of *Pseudomonas* (78–80).

## Materials and Methods

### Bacterial strains, plasmids and culture conditions

Chemicals were supplied by Sigma-Aldrich Co. (St. Louis, MO, USA) if not otherwise stated. All bacterial strains used in this study are listed in **Table S1**. Cultures of *P. putida* KT2440, *E. coli* and their derivatives were incubated at 30°C and 37°C, respectively. For standard applications, routine cloning procedures and during genome engineering manipulations, cells were grown in LB medium (containing 10 g L^−1^ tryptone, 5 g L^−1^ yeast extract and 10 g L^−1^ NaCl, pH = 7.0). All liquid cultures were agitated at 250 rpm (*E. coli*) or 180 rpm (*P. putida*) in an orbital MaxQ™ 8000 shaker incubator (ThermoFisher Scientific, Waltham, MA, USA). Solid media contained 15 g L^−1^ agar. Kanamycin (Km) and gentamicin (Gm) were added whenever needed to retain plasmids at 50 μg mL^−1^ and 10 μg mL^−1^, respectively. M9 minimal medium (21) supplemented with 20 mM citrate, 20 mM glucose, 24 mM ribose, 20 mM fructose, 40 mM glycerol or 20 mM gluconate was used for phenotypic characterization of *P. putida* KT2440 (note that the total amount of carbon atoms was kept constant across experimental conditions). Growth rates were determined fitting the temporal changes in optical density measured at 600 nm (OD_600_) to the exponential growth model. Optical densities were recorded in a microplate reader (Elx808, BioTek Instruments; Winooski, VT, USA).

For recording relative fluorescence intensity in the translation reporter constructs, cultures were grown in M9 media supplemented with the corresponding carbon source overnight. These cultures were used to inoculate wells of a 96-well-plate, containing the same media. The cultures were incubated shaken at 30°C in a Synergy HI plate reader (BioTek Instruments) and growth and fluorescence were followed by spectrophotometry at 600 nm and 500/530 nm (excitation/emission wavelength), respectively. The highest relative fluorescence intensity is reported, corresponding to the beginning of the stationary phase.

### General cloning procedures and plasmid construction

All oligonucleotides and plasmids used in this work are listed in **Table S1** and **Table S2**, respectively. Phusion™ Hot Start II high-fidelity DNA polymerases (ThermoFisher Scientific; Waltham, MA, USA) was used for DNA amplification according to the manufacturer’s specifications. For colony PCR, the commercial *OneTaq*™ master mix (New England BioLabs; Ipswich, MA, USA) was used according to the manufacturer’s instructions. *E. coli* DH5α was used for general cloning purposes, while *E. coli* DH5α λ*pir* was employed when cloning and maintaining replicons with the conditional, Π-dependent origin of replication *RK6* (**Table S2**). Chemically-competent *E. coli* cells were prepared and transformed with plasmids according to well-established protocols (81). *P. putida* was rendered electrocompetent by washing the biomass from saturated (24 h) LB medium cultures with 0.3 M sucrose, and cells were routinely transformed with plasmids by electroporation, following the protocol of Choi et al. (82).

Genomic DNA was extracted from *P. putida* KT2440 using the DNeasy Blood & Tissue Kit (Qiagen). Plasmids for gene deletion were constructed by amplification of 500-bp-long fragments up- and downstream of the corresponding gene. These fragments were spliced by overlap extension PCR and ligated into the digested backbones. USER cloning was performed according to (25). Similarly, the plasmids containing the promotor regions of *zwf* and *zwfA* were constructed by amplifying the corresponding genomic regions with the primer pairs P_*zwfA*__*Eco*RI-F/P_*zwfA*__*Bam*HI-R and P_*zwfB*__*Eco*RI-F/P_*zwfB*__*Bam*HI-R (**Table S1**), respectively, digesting the fragments and vector pSEVA237Y with *Eco*RI and *Bam*HI, followed by ligation. Integrity of all constructs was checked by DNA sequencing. Genomic DNA was employed as the template for the amplification of *zwfB* and *zwfC* with the primer pairs Zwf_*Nde*I-F/Zwf_*Bam*HI-R and ZwfB_*Nde*I-F/ZwfB_*Bam*HI-R (**Table S1**), respectively. Plasmid pET28a and the amplification products were restricted with *Nde*I and *Bam*HI, separated on a 1% (w/v) agarose gel and ligated. Ligation was accomplished using T4 DNA ligase (New England Biolabs), according to the protocol provided by the manufacturer and giving rise to expression plasmids pET28a·*zwfB^Pp^* and pET28a·*zwfC^Pp^*. Then, electrocompetent *E. coli* DH5α cells were transformed with these ligation products. Positive clones, identified by colony PCR, were verified by DNA sequencing.

### G6PDH activities in cell-free extracts

A 10-mL aliquot of M9 medium supplemented with the corresponding carbon source was inoculated with *P. putida* KT2440 cells from a single colony and incubated for 16 h. This culture was used to inoculate a 100-mL Erlenmeyer flask containing 20 mL of fresh medium to achieve an initial OD_600_ of 0.05. The culture was grown in an orbital shaker (180 rpm) to an OD_600_ of 0.5. Cells were then harvested (4000×*g*, 10 min, 4°C), the supernatant discarded and the pellet placed in an ice bath. The pellet was suspended in 0.5 mL of buffer A, containing 50 mM Tris·HCl, 5 mM NaCl, 5 mM MgCl_2_ and 5 % (v/v) glycerol, pH = 8.0. Cells were mixed with 0.3 g of glass beads (212-300 μm, acid washed, Sigma-Aldrich Co. G1277) and disrupted at 6,000 rpm for 20 s (Precellys24, Berlin Technology, Berlin, Germany). Unbroken cells and cell debris were pelleted by centrifugation at 17000×*g* for 2 min at 4°C. The supernatant was transferred to a new tube. Protein concentration of the supernatant was determined by means of the Bradford assay (83) using a commercial kit (Pierce, Thermo Fischer). For these measurements, 5 μg of total protein, 0.25 mM NAD(P)^+^ (as indicated in the text) and 2 mM G6P were mixed in 0.2 mL of buffer A, and incubated at 30°C. All stock solutions were freshly prepared. The initial rate of formation of NAD(P)H was followed by spectrophotometry at 340 nm in a Synergy HI plate reader (BioTek Instruments).

### Enzyme production and purification and kinetic assays

*E. coli* BL21(DE3) transformed either with plasmid pET28a·*zwfB* or pET28a·*zwfC* was grown in LB medium at 37°C with agitation until OD_600_ ≈ 0.5. At that point, isopropyl-β-D-1-thiogalactopyranoside (IPTG) was added to the cultures and the temperature of the incubator was decreased to 25°C. Cells were incubated for 16-20 h under these conditions. Cells from these IPTG-induced cultures were harvested by centrifugation (5000×*g*, 4°C, 30 min), washed twice with ice-cold buffer A and re-suspended in buffer A supplemented with 2 mM (L+D)- dithiothreitol, additional NaCl (up to 100 mM), 20 mM imidazole and a protease inhibitor cocktail (Roche) prepared as recommended by the manufacturer. Re-suspended cells were sonicated on ice and ultra-centrifuged (30,000×*g*, 4°C, 60 min). The supernatants obtained after this step were loaded in 5-mL His-trap columns (GE Healthcare Systems, Chicago, IL, USA) pre-equilibrated with buffer A supplemented with 100 mM NaCl and 20 mM imidazole. His-tagged proteins were eluted by gradually increasing the concentration of imidazole in the buffer flowing through the His-trap columns. The gradients of imidazole-containing buffer were linear (from 20 mM to 500 mM), corresponding to 40 times the His-trap column volume. Eluted fractions (1.5 mL) were collected and the G6PDH activity was measured by following the formation of NADPH by spectrophotometry at 340 nm as explained in the previous section. Fractions containing G6PDH activity levels in the upper quintile of all samples were pooled, concentrated and equilibrated in buffer A. Purity of G6PDH-B and G6PDH-C was evaluated by separation in an SDS-PAGE. Pure proteins were preserved with 50% (v/v) glycerol, added gradually to avoid precipitation of the purified enzyme preparation, and protein stocks were stored at –20°C until they were employed for kinetic assays.

Samples of pure protein were equilibrated in buffer A at 4°C before the kinetic assays. The specific activities of samples were compared with the values registered before the storage, and no significant decline in the activities was observed after 2 weeks of storage in the conditions mentioned above. The stocks of G6P, NAD and NADP were neutralized and their concentrations were determined by titration as described by Olavarría et al. (45). The kinetic assays were performed in buffer A, at 30°C, supplemented with NAD(P)^+^ and G6P at different concentrations as indicated in the text. An UV/Vis Synergy 2 spectrophotometer (BioTek Instruments) and non-binding flat-bottom 96-wells plates (model 655901, Greiner Bio-One GbmH, Kremsmünster, Austria) were used for recording spectrophotometric changes in the reaction mixtures. Previous assays were performed to determine the optimal concentration of enzyme to minimize enzyme inactivation **(Fig. S6)** and to obtain initial rate estimations before consumption of 5% of the initial amount of substrate (84). Such preliminary assays were also employed to obtain *a priori* estimations of K_M_ values using the method of linear direct plotting (85). Initial rates were estimated combining NAD(P)^+^ concentrations ranging from 20 to 2,000 μM with different fixed concentrations of G6P, ranging from 125 to 2,200 μM. Kinetic parameters obtained through the global fitting procedure were employed to estimate the relative formation of NADH and NADPH in the reactions catalyzed by G6PDH-B and G6PDH-C, according to a method previously described (19). The *DYNAFIT* software package (86) was used for this analysis. It allows for global fittings, i.e. using data obtained using different concentrations of substrate and/or enzyme and/or modifiers simultaneously.

### Phylogenetic analysis

Amino acid sequence of proteins with annotated G6PDH function were acquired from the orthologue database OrthoDB (57). Sequences of each taxonomic order were aligned (87) and categorized according to the key catalytic residue corresponding to R50 in *E. coli* (53). Proteins containing an arginine residue in the corresponding module of G6PDH were considered to be NADPH specific, while all other amino acids in this position were considered to have a loosened cofactor discrimination. In the following step, organisms carrying these variants were categorized as using (i) the ED pathway, if their genomes encoded the signature enzyme KDPG aldolase, or (ii) the EMP pathway, if they contained a fructose-6-phosphate-1-kinase annotated in the OrthoDB. Note that both KDPG aldolase and fructose-6-phosphate-1-kinase are both unique for each type of metabolic pathway (1). If none of these proteins were found, the predominant metabolism of the corresponding host was categorized as “unknown”. The phylogenetic tree, blending all these constraints, was drawn according to Jun et al. (88), and all sequence logos were generated with weblogo3 (89).

### Data and statistical analysis

Initial rates obtained in experiments with purified enzymes were globally fitted using the Dynafit software version 4 (Biokin Ltd., Watertown, MA, USA) (86). Beyond the estimation of the best fitted values for each kinetic parameter, this software enabled a model discrimination analysis among different reaction mechanisms to determine which of them best explained the observed results. Due to the non-linear relationship between rates and substrate concentrations, the uncertainty in the estimations of the kinetic parameters was expressed as 95% confidence intervals. All other experiments reported in this study were independently repeated at least in biological triplicates (as indicated in the corresponding figure or table legend), and the mean value of the corresponding parameter ± standard deviation is presented. Whenever relevant, the level of significance of the statistical differences was evaluated by means of the two-tailed, homoscedastic Student's *t* test, with α = 0.01 (**) or α = 0.05 (*), as indicated in the respective figure legends.

## Acknowledgments

This work was funded by The Novo Nordisk Foundation (individual grant NNF10CC1016517, and *LiFe*, NNF18OC0034818), the European Union’s *Horizon 2020* Research and Innovation Programme under grant agreement No. 814418 (*SinFonia*) and the Danish Council for Independent Research (*SWEET*, DFF-Research Project 8021-00039B) to P.I.N.

We declare that we have no competing financial interests.

## Supplemental Files

**Table S1.** Bacterial strains and plasmids used in this study.

**Table S2.** Oligonucleotides used in this study.

**Figure S1** | Confirmation of the genotypes of the *zwf* deletion strains.

**Figure S2** | Growth phenotypes of *zwf* deletion strains on M9 minimal medium.

**Figure S3** | Comparison of the genomic organization around *zwfC* in different species.

**Figure S4** | Transcriptional levels of *zwfA, zwfB* and *zwfC* in *P. putida* KT2440 growing on different carbon sources.

**Figure S5** | Alignment of the proteins encoded by *zwfA*, *zwfB* and *zwfC* in *P. putida* KT2440.

**Figure S6** | Selwyn plot of three representative reaction progress curves obtained with G6PDH-B at three different enzyme concentrations.

